# Cross-validation for the estimation of effect size generalizability in mass-univariate brain-wide association studies

**DOI:** 10.1101/2023.03.29.534696

**Authors:** Janik Goltermann, Nils R. Winter, Marius Gruber, Lukas Fisch, Maike Richter, Dominik Grotegerd, Katharina Dohm, Susanne Meinert, Elisabeth J. Leehr, Joscha Böhnlein, Anna Kraus, Katharina Thiel, Alexandra Winter, Kira Flinkenflügel, Ramona Leenings, Carlotta Barkhau, Jan Ernsting, Klaus Berger, Heike Minnerup, Benjamin Straube, Nina Alexander, Hamidreza Jamalabadi, Frederike Stein, Katharina Brosch, Adrian Wroblewski, Florian Thomas-Odenthal, Paula Usemann, Lea Teutenberg, Julia Pfarr, Andreas Jansen, Igor Nenadić, Tilo Kircher, Christian Gaser, Nils Opel, Tim Hahn, Udo Dannlowski

## Abstract

**Introduction:** Statistical effect sizes are systematically overestimated in small samples, leading to poor generalizability and replicability of findings in all areas of research. Due to the large number of variables, this is particularly problematic in neuroimaging research. While cross-validation is frequently used in multivariate machine learning approaches to assess model generalizability and replicability, the benefits for mass-univariate brain analysis are yet unclear. We investigated the impact of cross-validation on effect size estimation in univariate voxel-based brain-wide associations, using body mass index (BMI) as an exemplary predictor.

**Methods:** A total of n=3401 adults were pooled from three independent cohorts. Brain-wide associations between BMI and gray matter structure were tested using a standard linear mass-univariate voxel-based approach. First, a traditional non-cross-validated analysis was conducted to identify brain-wide effect sizes in the total sample (as an estimate of a realistic reference effect size). The impact of sample size (bootstrapped samples ranging from n=25 to n=3401) and cross-validation on effect size estimates was investigated across selected voxels with differing underlying effect sizes (including the brain-wide lowest effect size). Linear effects were estimated within training sets and then applied to unseen test set data, using 5-fold cross-validation. Resulting effect sizes (explained variance) were investigated.

**Results:** Analysis in the total sample (n=3401) without cross-validation yielded mainly negative correlations between BMI and gray matter density with a maximum effect size of *R*^2^_p_=.036 (peak voxel in the cerebellum). Effects were overestimated exponentially with decreasing sample size, with effect sizes up to *R*^2^_p_=.535 in samples of n=25 for the voxel with the brain-wide largest effect and up to *R*^2^_p_=.429 for the voxel with the brain-wide smallest effect. When applying cross-validation, linear effects estimated in small samples did not generalize to an independent test set. For the largest brain-wide effect a minimum sample size of n=100 was required to start generalizing (explained variance >0 in unseen data), while n=400 were needed for smaller effects of *R*^2^_p_ =.005 to generalize. For a voxel with an underlying null effect, linear effects found in non-cross-validated samples did not generalize to test sets even with the maximum sample size of n=3401. Effect size estimates obtained with and without cross-validation approached convergence in large samples.

**Discussion:** Cross-validation is a useful method to counteract the overestimation of effect size particularly in small samples and to assess the generalizability of effects. Train and test set effect sizes converge in large samples which likely reflects a good generalizability for models in such samples. While linear effects start generalizing to unseen data in samples of n>100 for large effect sizes, the generalization of smaller effects requires larger samples (n>400). Cross-validation should be applied in voxel-based mass-univariate analysis to foster accurate effect size estimation and improve replicability of neuroimaging findings. We provide open-source python code for this purpose (https://osf.io/cy7fp/?view_only=a10fd0ee7b914f50820b5265f65f0cdb).

## Introduction

Neuroimaging methods such as magnetic resonance imaging (MRI) have been used for several decades now to gain insights into the neurobiological underpinnings of psychological phenotypes. A plethora of scientific publications describes imaging-derived biomarkers of mental health disorders and perpetuates the hope to hope of utilizing such methods to aid clinical decision-making (Nour et al., 2022). However, recently the validity of findings involving relationships between neuroimaging and psychological phenotypes have been questioned due to accumulating reports of low replicability (Boekel et al., 2015; Genon et al., 2022; Marek et al., 2022). While such a replication crisis is not specific to neuroimaging research (e.g., for the field of psychology see Open Science Collaboration, 2015), its contributing factors may be particularly potent in this research domain due to high analytic flexibility in combination with a large number of statistical tests, which is likely to amplify publication bias (Botvinik-Nezer et al., 2020; Ioannidis, 2005; Jennings & Van Horn, 2012). Relatedly, replicability is undermined by small samples (Turner et al., 2018) which lead to unreliable and overestimated effect sizes (Button et al., 2013; Lane & Dunlap, 1978; Maxwell et al., 2008; Schönbrodt & Perugini, 2013). As studies particularly in neuroimaging research frequently use small samples (Elliott et al., 2020; Szucs & Ioannidis, 2020), this likely further contributes to low replicability. Recently, Marek et al. (2022) demonstrated that even without publication bias, associations between psychological and brain phenotypes are not replicable unless thousands of participants are included in the analysis. This seminal study has led to a vibrant discussion regarding replicability, sample size and effect size in the neuroimaging domain (Bandettini et al., 2022; Genon et al., 2022; Nour et al., 2022; Rosenberg & Finn, 2022; Spisak et al., 2023; Tervo-Clemmens et al., 2023). Notably, the demonstrated overestimation of effect sizes in small samples also occurs in statistically significant effects and, paradoxically, using more stringent significance thresholds even aggravates the problem (Lane & Dunlap, 1978; Marek et al., 2022).

In summary, the outlined evidence emphasizes an urgent need for solutions to increase the replicability of psychiatric neuroimaging findings. One possible solution that has been suggested repeatedly is to validate brain effects in independent data via cross-validation (Klapwijk et al., 2021; Kriegeskorte et al., 2010; Rosenberg & Finn, 2022). While cross-validation methods are standardly used in multivariate brain analyses to identify and counteract overfitting and assure generalizability of models (e.g., Redlich et al., 2014, 2016; Repple et al., 2023; Schaffer, 1993), they are rarely ever used on univariate brain effects, likely also because common neuroimaging analysis software packages do not offer options to conduct cross-validation. However, specifically in the domain of neuroimaging research, cross-validation methods may be useful even for univariate models as the outlined overestimation of effects can be seen as an overfitting of models in the face of high analytic flexibility in combination with a high number of tests in this domain. Thus, we investigate the utility of cross-validation for accurate estimation of effect size in univariate voxel-based brain analysis. Body mass index (BMI) is used as an exemplary predictor due to previous reports of good replicability of BMI with brain structure, as well as its pivotal relevance for various mental disorders (Bond et al., 2014; McWhinney et al., 2022; Opel et al., 2015, 2021). Due to reports of relatively large effect size estimates of the association between BMI and brain structure even in large samples (maximum effect size corresponding to approximately 2.7% explained variance; Opel et al., 2021), this predictor is suitable to investigate the impact of cross-validation for a broader range of effect size, as compared to other predictors where effect size estimates across brain modalities have been shown to be considerably smaller (Marek et al., 2022; Winter et al., 2022). In order to evaluate the relevance of cross-validation for generalizability of effects sizes as a function of the sample size, we conducted our analyses across different sample sizes.

In light of the current lack of brain analysis software implementations regarding this matter, we provide open-source Python code to enable other researchers to apply cross-validation for voxel-based brain analysis.

## Method

### Participants

A total of n=3401 participants were included from three independent German cohorts: the Marburg-Münster Affective Disorders Cohort Study (MACS; n=1655), the Münster Neuroimaging Cohort (MNC; n=722) and the BiDirect cohort (n=1024). All three cohorts include individuals with major depressive disorder (MDD) and healthy controls (HC) free from any lifetime mental disorder diagnoses. General study methods, exclusion criteria and cohort characteristics were comprehensively described elsewhere (MACS: Kircher et al., 2019 and Vogelbacher et al., 2018; MNC: Dannlowski et al., 2016 and Opel et al., 2019; BiDirect: Teismann et al., 2014). See supplements for an additional description of data exclusion steps and sample characteristics specific to the current analysis.

### Measures and procedure

BMI was calculated based on self-reported height and weight of participants. T1-weighted high-resolution anatomical brain images were acquired using 3T MRI scanner in all three studies. For the MACS sample two different MRI scanners were used at the recruitment sites in Marburg (Tim Trio, Siemens, Erlangen, Germany; combined with a 12-channel head matrix Rx-coil) and Münster (Prisma, Siemens, Erlangen, Germany; combined with a 20-channel head matrix Rx-coil). Data from the MNC and BiDirect samples were acquired using the same Gyroscan Intera scanner (later with Achieva update; Philips Medical Systems, Best, The Netherlands). Image preprocessing was conducted using the CAT12-toolbox (Gaser et al., 2022; https://neuro-jena.github.io/cat/) using default parameters for all samples. Briefly, images were bias-corrected, tissue classified, and normalized to MNI-space using linear (12-parameter affine) and non-linear transformations, within a unified model including high-dimensional geodesic shooting normalization (Ashburner & Friston, 2011). The modulated gray matter images were smoothed with a Gaussian kernel of 8 mm FWHM. Absolute threshold masking with a threshold value of 0.2 was used for all second level analyses as recommended for VBM analyses (https://neuro-jena.github.io/cat12-help/). Image quality was assessed by visual inspection as well as by using the check for homogeneity function implemented in the CAT12 toolbox.

### Statistical analysis

We investigated the association between BMI and gray matter brain structure using an established mass-univariate voxel-based morphometry (VBM) approach. To that end, BMI was used as a predictor in a general linear model (GLM) to predict voxel-wise gray matter density. The following nuisance parameters were included in the model: age, sex, total intracranial volume (TIV) and four dummy-coded scanner variables to control for scanner hardware differences. Two-sided whole brain effects of BMI were tested and partial *R*^2^ (*R*^2^_p_) was used as a measure of effect size. To assess the impact of sample size and cross-validation on the estimation of voxel-based effect sizes the following steps were undertaken (for a schematic overview see Figure 1):

**Figure 1.**
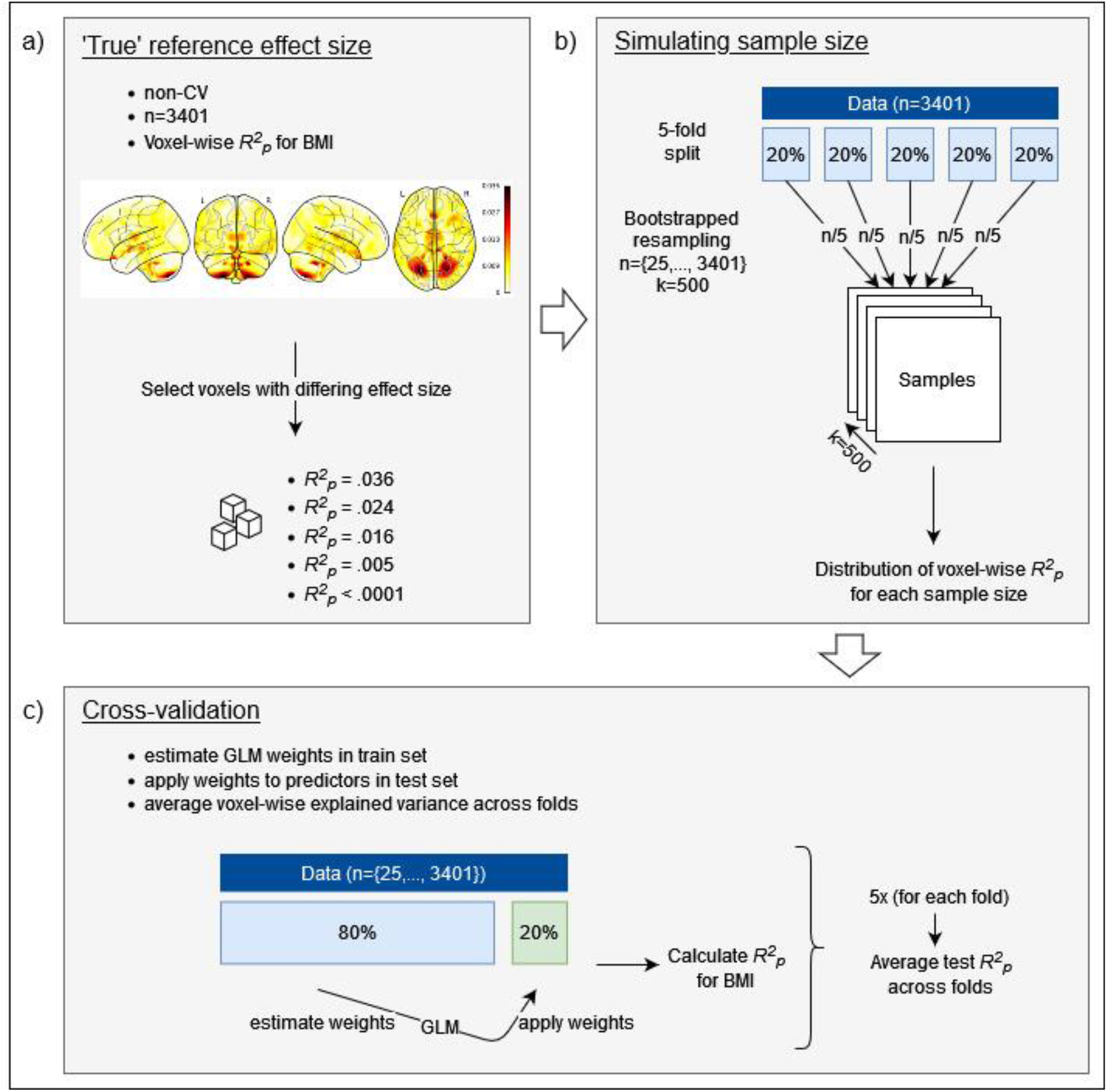
Analysis steps to obtain voxel-wise effect size estimates across sample sizes with and without cross-validation *Note. CV, cross-validation; BMI, Body Mass Index; GLM, general linear model*.

1. Classical non-cross-validated analysis was conducted in the maximum available sample without bootstrapping (n=3401). The resulting voxel-wise explained variance of BMI was used as an estimate for the realistic underlying effect size (in the following referred to as the *reference effect size*). Subsequently, voxels were selected covering a range of different representative effect sizes, including the voxel with the largest brain-wide reference effect size. In addition, the voxel with the smallest brain-wide reference effect size was selected to investigate effect size estimation based on an underlying null effect. An uncorrected significance threshold of p<.001 and extent threshold of k>200 was used for visualization purposes of brain-wide associations.
2. Based on the findings of Marek et al. (2022) we subsequently investigated the influence of sample size on the non-cross-validated effect sizes. Sample size was manipulated using samples of 18 different sizes: n=25, 35, 50, 70, 100, 150, 200, 300, 400, 600, 800, 1000, 1300, 1600, 2000, 2400, 2900, 3401. For each preselected voxel a bootstrapped resampling distribution of effect size was obtained containing k=500 bootstrap runs per sample size. For each sample size the mean effect size was calculated across all bootstrap runs, as well as a 95% confidence interval (CI).
3. Finally, the impact of cross-validation on effect size estimates in the selected voxels was investigated across samples created with the resampling procedure described above. To this end, 5-fold cross-validation was applied by estimating linear effects using the GLM described above within respective train sets and then applying the resulting linear coefficients to the respective test sets. To this end the resulting beta coefficients yielded by model estimation within the train set was applied to the design matrix (individual predictor values) of the test set (i.e., data new to the trained linear model). The predictive value of the BMI predictor was evaluated by calculating the explained variance (based on residuals) with and without BMI as a predictor in the model. Then the mean *R*^2^_p_ for BMI was calculated across the five test sets. This was used to quantify the extent to which the linear voxel-wise coefficients for BMI obtained from the train sets explain variance within unknown test data (i.e., generalization of effects to unseen data). Note that while *R*^2^_p_ usually has a range from 0-1, it can become negative (and even exceed −1) in this case due to the application of linear coefficients to unseen data which can result in an effect size lower than prediction by the mean within the same sample (normally the baseline reference corresponding to an effect size *R*^2^_p_ =0). A *point of initial generalizability* was defined as the minimum sample size necessary to achieve positive effect sizes in the test sets (i.e., the lower bound of the 95% CI above *R*^2^_p_ =0). This point of initial generalizability can be interpreted as the minimum sample size needed for linear effects of a given effect size to generalize to unknown samples in a 5-fold cross-validation framework. The 5-fold cross-validation was chosen over a higher number of folds to be able to simulate very small (but commonly used) samples sizes. In order investigate the impact of this cross-validation on effect size estimation across different sample sizes we applied the same bootstrap approach described above also to cross-validation-based effect size estimates, resulting in resampling distributions of average test set effect sizes for each sample size (95% CI were based on these bootstrap runs). To allow this combination of cross-validation and bootstrapping, we first conducted the 5-fold split of the total sample and then performed the bootstrapped resampling for each sample size by drawing with replacement from each split separately – thus only allowing replacement within one split but not across splits. This is necessary to avoid data leakage between train and test sets.

All analyses were conducted in Python (version 3.9.12) using the nilearn package (version 0.9.1) for voxel-based GLMs and k-Fold from the sklearn package (version 1.1.1) for cross-validation. The complete analysis code is provided online (https://osf.io/cy7fp/?view_only=a10fd0ee7b914f50820b5265f65f0cdb).

## Results

### Associations between BMI and gray matter – identification of reference effect sizes in n=3401

The association between BMI and gray matter density in a non-cross-validated standard analysis is shown in Figure 2a and supplementary Figure S1. Liberal thresholding at uncorrected p<.001, k>200 resulted in widespread clusters across the brain. Effect size in significant voxels ranged up to *R*^2^_p_ =.036. Peak voxel coordinates of significant clusters covering a wide range of effect sizes, as well as the voxel with the brain-wide smallest effect size were selected for further analysis (see Figure 2a). The selected voxels from largest to smallest effects were located in 1) the cerebellum (*R*^2^_p_=.036, p<.0001, x-24, y-72, z-60), 2) the posterior medial orbitofrontal cortex (mOFC; *R*^2^_p_ =.024, p<.0001, x-4, y25, z-30), 3) the thalamus (*R*^2^_p_=.016, p<.0001, x6, y-12, z10), 4) the anterior mOFC (*R*^2^_p_=.005, p<.0001, x4, y69, z-6) and 5) the calcarine (*R*^2^_p_ <.0001, p=.999, x-3, y-80, z12).

**Figure 2.**
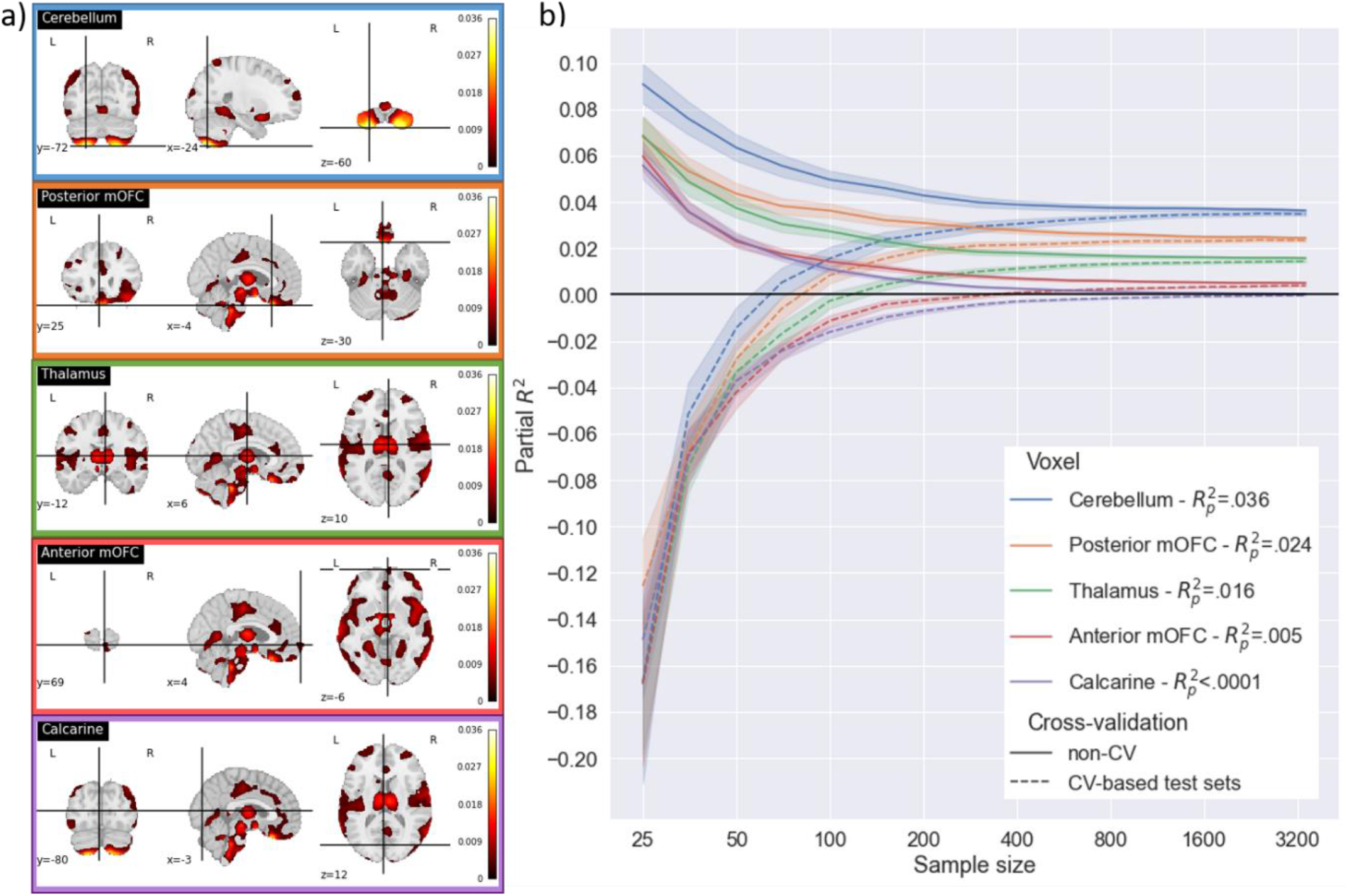
The association between Body Mass Index and gray matter density - selected voxel locations and their effect size estimates across sample sizes, with and without and cross-validation Note. a) Shows non-cross-validated two-sided effect of BMI on gray matter density at a liberal uncorrected threshold of p<.001 in the maximum sample of n=3401. The location of selected peak voxel coordinates in the cerebellum (largest brain-wide effect), posterior medial prefrontal cortex (mOFC), thalamus, anterior mOFC and calcarine (smallest brain-wide effect) are shown. Color bars represent R^2^_p_ values. b) Shows effect size estimates for the five different voxels across sample sizes (resampling with k=500 per sample size). Non-cross-validated (non-CV) effect sizes are presented with solid lines and test set effect sizes with dotted lines. Error bands represent the 95% confidence interval. Respective effect sizes of selected voxels in non-cross-validated analysis of full sample are presented in the legend (ranging from η^2^_p_=.036 to η^2^_p_<.0001).

### The impact of sample size on non-cross-validated effect size estimation

Effect size was systematically overestimated in small samples in the non-cross-validated analysis. Averaged across all bootstrapped samples with size n=25, effect size was inflated approximately .06 *R*^2^_p_ units for all five voxels, resulting in a 2.5-fold inflation for the voxel with the largest reference effect size (*R*^2^_p_ =.091 instead of *R*^2^_p_ =.036) and 11.9-fold inflation for the voxel with a reference effect size of *R*^2^_p_ =.005. Maximum effect size estimates went as high as *R*^2^_p_ =.734 in samples of n=25. Even the voxel with the brain-wide lowest reference effect size (null-effect) reached a maximum effect size of up to *R*^2^ =.429 (average *R*^2^_p_ =.056) in these samples with n=25. Average estimates of effect size (as well as maximum estimates) decreased exponentially with increasing sample size.

Inspection of partial correlation coefficients showed that a broad range of associations was found across bootstrapped samples, particularly with small sample sizes. For the voxel with the largest brain-wide reference effect size, associations ranging from a negative correlation r=-.73 to a positive correlation r=.60 were found in samples with n=25 (see supplementary Figure S2).

Detailed summary statistics for effect size estimates across sample sizes and voxels are presented in supplementary Tables S2-6. While the average effect size estimate across bootstrapped samples may be informative, it should be noted that the distribution of estimates was highly skewed with some extreme outliers. This distribution across single runs from the resampling procedure is further visualized in supplementary Figure S3.

### The impact of cross-validation on effect size estimation

On average test set effect size estimates were descriptively lower compared to non-cross-validated effect sizes across all sample sizes and all voxels (this was true for 93.83% of bootstrapped samples across all sample sizes). This disparity was particularly strong in small samples where non-cross-validated effect sizes were largest while test set effect sizes were mostly negative (indicating no generalization of linear effects to unknown samples).

A *point of initial generalizability* was reached after n=100 for the largest reference effect size, while larger samples were needed to reach this point for voxels with smaller effect size: n=100 for *R*^2^_p_ =.024, n=150 for *R*^2^_p_ =.016, and n=400 for *R*^2^_p_ =.005. For the voxel with the minimum brain-wide effect size this *point of initial generalizability* was never reached, as expectable for an underlying null-effect. In fact, for this voxel even the mean and upper bound of the 95% CI were always below an effect size of zero at any sample size, when applying cross-validation, indicating effective protection against false-positives (see supplementary Tables S2-6).

Estimates derived from non-cross-validated analysis and from test sets using cross-validation approached convergence in larger samples. In samples with n=3401, non-cross-validated and test set effect sizes differed only marginally (average difference across voxels: *R*^2^_p_ =.001). However, full convergence was never reached between non-cross-validated and test set effect size estimates, meaning that confidence intervals did not overlap, even at largest sample sizes. Detailed descriptive statistics for effect size estimates across sample sizes, voxels and cross-validation are given in supplementary Tables S2-6.

## Discussion

Using BMI as an exemplary predictor variable, we investigated the utility of cross-validation for the accurate estimation of effect sizes of univariate brain-wide associations. Replicating previous findings (Marek et al., 2022), we find that effect size estimates are highly overestimated and unreliable in small samples if no cross-validation is applied. Furthermore, we demonstrate that cross-validation can be used to reveal poor generalizability to new data of these overestimated effects in small samples (indicated by negative effect size estimates). In larger samples, cross-validation-derived test set effect sizes start becoming positive, suggesting a *point of initial generalizability* that can be interpreted as the minimum sample size necessary to achieve generalizable linear effect estimates.

Importantly, sample sizes of several hundreds of participants may be sufficient to accurately estimate brain-wide associations, given sufficiently large underlying effect sizes. The cross-validation approach could facilitate a differentiation between artificially inflated large effects and robust ‘true’ large effects. The standardly implementation of cross-validation to assess the generalizability of brain-wide effects should be considered in addition to conventional reporting of significance and traditional effect size estimates. The utilization of cross-validation to facilitate generalizability of neuroimaging findings has been repeatedly suggested (Klapwijk et al., 2021; Kriegeskorte et al., 2010; Rosenberg & Finn, 2022). However, to the best of our knowledge, we are the first to systematically investigate the utility of cross-validation for effect size estimation in brain-wide univariate analysis.

The presented overestimation of effect sizes in small samples falls in line with previous findings in the literature (Button et al., 2013; Lane & Dunlap, 1978; Maxwell et al., 2008; Schönbrodt & Perugini, 2013). Marek et al. (2022) demonstrated that even the largest brain-wide associations of compound cognitive and psychopathological variables with brain structure and function corresponded to less than 0.5% explained variance, which was inflated to up to approximately 40% explained variance in small samples. The authors conclude that thousands of individuals are needed for robust and replicable estimates in this research domain. While these findings are striking and important, the authors’ conclusion has been questioned by various scholars. The main critique is that true effect sizes may be considerably higher under ideal conditions leading to smaller sample sizes necessary for accurate estimation of effects (DeYoung et al., 2022). In a study of over 1800 adults, Winter et al. (2022) investigated univariate brain-wide, cross-modal brain differences between HC and lifetime MDD individuals and conclude that largest effect sizes go up to 1.7% explained variance. However, they further report that comparisons of HC with chronic or acute MDD individuals yield higher effects sizes of up to 2.7% explained variance (in n>900). Further, studies suggest that effect sizes are larger in other diagnostic groups such as psychotic disorders (Hettwer et al., 2022). Similarly, Repple and Gruber et al. (2022) present transdiagnostic structural connectome alterations, with largest effect sizes found in patients with schizophrenia, possibly rendering smaller samples sufficient to detect effects.

Our findings expand the results by Marek et al. (2022) to a voxel-based analysis framework and to a broader range of effect sizes. Importantly, we provide evidence that for larger effect sizes – which can occur under specific conditions as outlined above – hundreds of participants could suffice to accurately estimate linear brain-wide effects. When smaller effect sizes are assumed, our findings are highly comparable to the findings by Marek et al. as we similarly find that a ‘true’ effect size of approximately ∼0.5% explained variance requires very large samples to obtain robust estimates of effects, although cross-validation may enable the robust detection of accurate estimates already in somewhat smaller samples (n>400). The ongoing debate surrounding effect size and sample size in the neuroimaging domain stresses the importance of consistent reporting of effect sizes in publications, as well as interpreting effect sizes in the context of sample size.

While the above considerations could guide sample size planning of neuroimaging studies, a problem arises when the ‘true’ effect size is unknown (and effect size inflations in small samples make it particularly difficult to estimate this solely from existing literature). How can researchers know if an obtained effect size is accurate or inflated? Using a cross-validation framework, we demonstrate that the application of identified linear effects to unseen data can be utilized to identify inflated, non-generalizable effect sizes. Notably, this procedure can be applied in smaller studies to verify if a putative large effect size is robust and may warrant the use of a smaller sample. Importantly, our findings are in line with low test set performance merely reflecting an underpowered sample and not necessarily meaning that an effect does not exist (Helwegen et al., 2023). In other words, even substantial effects of 3.6% explained variance can barely be differentiated from null effects in small samples based on cross-validation-derived effect sizes alone. However, the relative congruence between non-cross-validated and cross-validation-derived effect sizes can provide a strong argument for robust and replicable linear associations with realistic effect size estimates. Thus, calculating non-cross-validated and cross-validation-derived effect sizes combined with a thorough inspection of the similarity between the resulting estimates could be informative. Notably, while cross-validation decreases the probability of effect size inflation, it does not eliminate it. Even unrealistically high effect size estimates can in rare cases generalize to unseen test set data, particularly in small samples, as shown by some extreme outliers in our results.

While it is difficult to deduct a clear recommendation for sample sizes based on our cross-validation analyses, we propose that a *point of initial generalizability* can be defined as the sample size where linear effects start explaining variance in unseen data. This aspect could expand the traditional power analysis framework by the question of generalizability in addition to significance. In other words, while traditional power analysis answers the question of how large a sample needs to be for a given effect to become *significant*, our results open up the possibility for defining sample sizes necessary for linear effects to *generalize* to unseen data.

In large samples effect sizes became highly stable and barely differed whether cross-validation was applied or not. This finding implicates that if several thousands of individuals are available (e.g., due to consortia data), within sample k-fold cross-validation may barely alter the result, regardless of the underlying true effect size.

The current study entails several important limitations. Firstly, it is unclear whether our findings can be generalized to other cross-validation methods. While 5-fold cross-validation was chosen for the current analysis so that small samples of n=25 could be included, other splits (e.g., 10-fold) could in principle also be suitable to be applied in voxel-based univariate brain analysis. While a systematic comparison of different cross-validation methods for the application to univariate voxel-based analysis is beyond the scope of this work, this may be a fruitful task for future studies. Further, it is open for discussion to what extent our findings are generalizable to other MRI modalities, such as functional MRI and parcellation-based brain analysis, as well as to predictors of other domains. We believe that generalizability to other voxel-based imaging modalities and other predictors is given, as the general pattern of results should not be specific to VBM effects of BMI. Parcellation-based brain analysis approaches could also profit from cross-validation. However, cross-validation could possibly be particularly beneficial in settings that are at higher risk for overfitting statistical models. Such risk is likely to be higher in more complex models, and in settings with higher analytical flexibility and a higher number of statistical tests (the latter being higher in voxel-based as compared to parcellation-based analysis). Interestingly, cross-validation does not seem to ultimately protect from an inflation of effect size in all settings. It has been shown that in complex multivariate modelling of voxel-based associations with psychopathology, even cross-validation-based performance measures are inflated in underpowered studies (Flint et al., 2021). In the same vein, we demonstrated that even cross-validation-derived effect sizes can be inflated in small samples although the probability of an overestimation of effects is lower as compared to non-cross-validated estimates (as outlined above).

In summary the utilization of cross-validation can contribute two major benefits: 1) identify if a study is underpowered and corresponding effect size estimates are likely to be highly inflated and 2) corroborate the accuracy of an effect size estimate of a sufficiently powered study (also for underlying null-effects). Thus, we propose that cross-validation procedures should be applied to foster replicability of neuroimaging research and facilitate accurate estimation of effect sizes. Further, we provide concrete steps for an application of cross-validation to mass-univariate voxel-based analysis, as well as corresponding open-source Python code.

## Supporting information

Supplementary methods and results

## Acknowledgements

This work is part of the German multicenter consortium “Neurobiology of Affective Disorders. A translational perspective on brain structure and function”, funded by the German Research Foundation (Deutsche Forschungsgemeinschaft DFG; Forschungsgruppe/Research Unit FOR2107).

Principal investigators (PIs) with respective areas of responsibility in the FOR2107 consortium are: Work Package WP1, FOR2107/MACS cohort and brainimaging: Tilo Kircher (speaker FOR2107; DFG grant numbers KI 588/14-1, KI 588/14-2), Udo Dannlowski (co-speaker FOR2107; DA 1151/5-1, DA 1151/5-2), Axel Krug (KR 3822/5-1, KR 3822/7-2), Igor Nenadic (NE 2254/1-2), Carsten Konrad (KO 4291/3-1). CP1, biobank: Petra Pfefferle (PF 784/1-1, PF 784/1-2), Harald Renz (RE 737/20-1, 737/20-2). CP2, administration. Tilo Kircher (KI 588/15-1, KI 588/17-1), Udo Dannlowski (DA 1151/6-1), Data access and responsibility: All PIs take responsibility for the integrity of the respective study data and their components. All authors and coauthors had full access to all study data.

Acknowledgements and members by Work Package (WP): WP1: Henrike Bröhl, Katharina Brosch, Bruno Dietsche, Rozbeh Elahi, Jennifer Engelen, Sabine Fischer, Jessica Heinen, Svenja Klingel, Felicitas Meier, Tina Meller, Torsten Sauder, Simon Schmitt, Frederike Stein, Annette Tittmar, Dilara Yüksel (Dept. of Psychiatry, Marburg University). Mechthild Wallnig, Rita Werner (Core-Facility Brainimaging, Marburg University). Carmen Schade-Brittinger, Maik Hahmann (Coordinating Centre for Clinical Trials, Marburg). Michael Putzke (Psychiatric Hospital, Friedberg). Rolf Speier, Lutz Lenhard (Psychiatric Hospital, Haina). Birgit Köhnlein (Psychiatric Practice, Marburg). Peter Wulf, Jürgen Kleebach, Achim Becker (Psychiatric Hospital Hephata, Schwalmstadt-Treysa). Ruth Bär (Care facility Bischoff, Neunkirchen). Matthias Müller, Michael Franz, Siegfried Scharmann, Anja Haag, Kristina Spenner, Ulrich Ohlenschläger (Psychiatric Hospital Vitos, Marburg). Matthias Müller, Michael Franz, Bernd Kundermann (Psychiatric Hospital Vitos, Gießen). Christian Bürger, Katharina Dohm, Fanni Dzvonyar, Verena Enneking, Stella Fingas, Katharina Förster, Janik Goltermann, Dominik Grotegerd, Hannah Lemke, Susanne Meinert, Nils Opel, Ronny Redlich, Jonathan Repple, Kordula Vorspohl, Bettina Walden, Dario Zaremba (Dept. of Psychiatry, University of Münster). Harald Kugel, Jochen Bauer, Walter Heindel, Birgit Vahrenkamp (Dept. of Clinical Radiology, University of Münster). Gereon Heuft, Gudrun Schneider (Dept. of Psychosomatics and Psychotherapy, University of Münster). Thomas Reker (LWL-Hospital Münster). Gisela Bartling (IPP Münster). Ulrike Buhlmann (Dept. of Clinical Psychology, University of Münster).

CP1: Julian Glandorf, Fabian Kormann, Arif Alkan, Fatana Wedi, Lea Henning, Alena Renker, Karina Schneider, Elisabeth Folwarczny, Dana Stenzel, Kai Wenk, Felix Picard, Alexandra Fischer, Sandra Blumenau, Beate Kleb, Doris Finholdt, Elisabeth Kinder, Tamara Wüst, Elvira Przypadlo, Corinna Brehm (Comprehensive Biomaterial Bank Marburg, Marburg University). Supplementary information is available at MP’s website.

