## Supplementary methods and results for "Cross-validation for the estimation of effect size generalizability in mass-univariate brain-wide association studies"

### Supplements

#### Supplementary Tables

Table S1. Sample characteristics across cohorts.

Table S2. Effect size estimates for **cerebellum** voxel with ‘true’ effect size of $R^2_p$**=.036** across sample size and cross-validation.

Table S3. Effect size estimates for **posterior mOFC** voxel with ‘true’ effect size of $R^2_p$**=.024** across sample size and cross-validation.

Table S4. Effect size estimates for **posterior mOFC** voxel with ‘true’ effect size of $R^2_p$**=.024** across sample size and cross-validation.

Table S5. Effect size estimates for **anterior mOFC** voxel with ‘true’ effect size of $R^2_p$**=.005** across sample size and cross-validation.

Table S6. Effect size estimates for **Calcarine** voxel with ‘true’ effect size of $R^2_p$**<.0001** across sample size and cross-validation.

#### Supplementary Figures

Figure S1. Non-thresholded brain-wide partial correlation between BMI and gray matter density in full sample (n=3401) without cross-validation.

Figure S2. Distribution of **partial *r*-coefficients** for **non-cross-validated** linear association between Body Mass Index and voxel-based morphometry across voxels and samples sizes.

Figure S3. Distribution of **partial $R^2$** estimates for **non-cross-validated** linear association between Body Mass Index and voxel-based morphometry across voxels and samples sizes.

Figure S4. Distribution of **test-set partial $R^2$** estimates for **cross-validated** linear association between Body Mass Index and voxel-based morphometry across voxels and samples sizes.

Figure S5. Association between non-cross-validated and cross-validation-based test set effect size estimates

#### Supplementary Methods

*Cohort characteristics and inclusion criteria*

The existing Marburg-Münster Affective Disorders Cohort Study (MACS), Münster Neuroimaging Cohort (MNC) and BiDirect study were accessed for analysis. The cohorts include adult participants with age 18-65 years (35-65 years for the BiDirect study) and were all created to investigate affective disorders. Largest groups within each of the three cohorts comprise healthy control participants (HC) free from any mental disorder and participants with a major depressive disorder (MDD) diagnosis. Additional smaller patient groups with bipolar disorder (MACS and MNC), psychosis spectrum disorders (schizoaffective and schizophrenia; MACS), as well as comorbid cardiovascular diseases (BiDirect) were specifically recruited for the cohorts, and psychiatric comorbidities were generally admitted. MACS and BiDirect recruited patient populations from psychiatric hospitals and outpatient centers in and around Münster, Germany (MACS and BiDirect) and Marburg, Germany (MACS). For the MNC, inpatient populations were included only, recruited also at psychiatric hospitals in Münster, Germany.

*Inclusion criteria for current analysis*

Identical inclusion criteria were applied for all three independent cohorts. The following participants were excluded to create the analysis sample: 1) duplicate cases resulting from individuals that were included in more than one of the utilized cohorts, 2) all participants with bipolar disorder, psychosis spectrum disorder, substance dependencies, 3) all participants with severe head trauma, severe/chronic somatic illness (e.g., parkinson's disease, multiple sclerosis, stroke, myocardial infarction), 4) participants with a non-caucasian decent, 5) participants with imaging dropouts, such as missing MRI data and image artefacts, 5) participants with missing data in the Body Mass Index variable.

Table S1. Sample characteristics across cohorts.

| | MACS (n=1655) | | | MNC (n=722) | | | BiDirect (n=1024) | | |
|---|---|---|---|---|---|---|---|---|---|
| | Mean | SD | Count | Mean | SD | Count | Mean | SD | Count |
| Age | 35.35 | 13.05 | - | 36.91 | 12.2 | - | 50.58 | 7.79 | - |
| Sex (f/m) | - | - | 1074/581 | - | - | 376/346 | - | - | 558/466 |
| Diagnosis (HC/MDD) | - | - | 863/792 | - | - | 479/243 | - | - | 472/552 |
| BMI | 25.07 | 5.23 | - | 25.09 | 4.47 | - | 27.39 | 5.06 | - |

*Note. MACS, Marburg-Münster Affective Disorders Cohort Study; MNC, Münster Neuroimaging Cohort; SD, standard deviation; f, female; m, male; HC, health control participants; MDD, major depressive disorder participants; BMI, body mass index.*

Table S2. Effect size estimates for **cerebellum** voxel with reference effect size of $\boldsymbol{R^2_p}$**=.036** across sample size and cross-validation.

| Sample size | $R^2_p$ non-CV | | | | | CV-based $R^2_p$ test | | | | |
|---|---|---|---|---|---|---|---|---|---|---|
| | Mean | SD | 95% CI | Min | Max | Mean | SD | 95% CI | Min | Max |
| 25 | 0.0908 | 0.1001 | [0.082, 0.0996] | 0 | 0.5345 | -0.1669 | 0.4691 | [-0.2082, -0.1257] | -6.756 | 0.6347 |
| 35 | 0.076 | 0.0835 | [0.0686, 0.0833] | 0 | 0.4648 | -0.0517 | 0.1525 | [-0.0651, -0.0383] | -1.046 | 0.4698 |
| 50 | 0.0634 | 0.0678 | [0.0574, 0.0693] | 0 | 0.4019 | -0.0144 | 0.104 | [-0.0235, -0.0052] | -0.8571 | 0.3244 |
| 70 | 0.0556 | 0.0539 | [0.0509, 0.0603] | 0 | 0.307 | 0.0052 | 0.0717 | [-0.0011, 0.0115] | -0.3098 | 0.2401 |
| 100 | 0.0496 | 0.0445 | [0.0457, 0.0535] | 0 | 0.2273 | 0.0156 | 0.056 | [0.0107, 0.0205] | -0.2471 | 0.2012 |
| 150 | 0.046 | 0.0348 | [0.0429, 0.049] | 0 | 0.1886 | 0.0238 | 0.038 | [0.0204, 0.0271] | -0.0915 | 0.1836 |
| 200 | 0.0428 | 0.0302 | [0.0401, 0.0454] | 0 | 0.1831 | 0.0261 | 0.0322 | [0.0233, 0.029] | -0.053 | 0.1806 |
| 300 | 0.0398 | 0.0239 | [0.0377, 0.0419] | 0.0001 | 0.1283 | 0.0294 | 0.0256 | [0.0271, 0.0316] | -0.0331 | 0.125 |
| 400 | 0.0388 | 0.0208 | [0.037, 0.0406] | 0.0007 | 0.1163 | 0.0305 | 0.0218 | [0.0286, 0.0324] | -0.0212 | 0.1052 |
| 600 | 0.0379 | 0.0171 | [0.0364, 0.0394] | 0.0047 | 0.1021 | 0.0322 | 0.0174 | [0.0307, 0.0338] | -0.0074 | 0.0971 |
| 800 | 0.0374 | 0.0146 | [0.0361, 0.0387] | 0.0076 | 0.0884 | 0.0331 | 0.0149 | [0.0317, 0.0344] | -0.0038 | 0.0851 |
| 1000 | 0.0372 | 0.0129 | [0.0361, 0.0384] | 0.009 | 0.0823 | 0.0337 | 0.013 | [0.0326, 0.0349] | 0.002 | 0.08 |
| 1300 | 0.0372 | 0.0113 | [0.0362, 0.0382] | 0.0159 | 0.0806 | 0.0343 | 0.0116 | [0.0333, 0.0354] | 0.0118 | 0.0738 |
| 1600 | 0.0371 | 0.0103 | [0.0361, 0.038] | 0.0159 | 0.0762 | 0.0346 | 0.0105 | [0.0337, 0.0355] | 0.0128 | 0.0724 |
| 2000 | 0.0368 | 0.0088 | [0.036, 0.0376] | 0.0156 | 0.0651 | 0.0347 | 0.009 | [0.0339, 0.0355] | 0.0127 | 0.0621 |
| 2400 | 0.0367 | 0.0085 | [0.036, 0.0375] | 0.014 | 0.067 | 0.0349 | 0.0086 | [0.0342, 0.0357] | 0.0124 | 0.0644 |
| 2900 | 0.0366 | 0.0075 | [0.0359, 0.0372] | 0.0185 | 0.0647 | 0.0349 | 0.0075 | [0.0342, 0.0356] | 0.0176 | 0.0635 |
| 3401 | 0.0362 | 0.0068 | [0.0356, 0.0368] | 0.0189 | 0.0607 | 0.0347 | 0.0068 | [0.0341, 0.0353] | 0.0156 | 0.0601 |

*Note*. Effect size estimates are shown for non-cross-validated analysis (non-CV) and for the test sets of cross-validated analysis (model generalization to unknown data). Presented statistics are summarized across k=500 bootstrapped resamples of the respective sample size. mOFC, medial orbitofrontal cortex; SD, standard deviation; CI, confidence interval.

Table S3. Effect size estimates for **posterior mOFC** voxel with reference effect size of $\boldsymbol{R^2_p}$**=.024** across sample size and cross-validation.

| Sample size | $R^2_p$ non-CV | | | | | CV-based $R^2_p$ test | | | | |
|---|---|---|---|---|---|---|---|---|---|---|
| | Mean | SD | 95% CI | Min | Max | Mean | SD | 95% CI | Min | Max |
| 25 | 0.068 | 0.0867 | [0.0604, 0.0756] | 0 | 0.5843 | -0.1258 | 0.2329 | [-0.1462, -0.1053] | -2.3819 | 0.4661 |
| 35 | 0.0534 | 0.067 | [0.0475, 0.0593] | 0 | 0.4646 | -0.068 | 0.1503 | [-0.0812, -0.0548] | -1.0998 | 0.4718 |
| 50 | 0.0435 | 0.0496 | [0.0392, 0.0479] | 0 | 0.2859 | -0.0279 | 0.0873 | [-0.0356, -0.0202] | -0.8283 | 0.2631 |
| 70 | 0.0382 | 0.0404 | [0.0346, 0.0417] | 0 | 0.2554 | -0.006 | 0.0544 | [-0.0108, -0.0013] | -0.2514 | 0.2046 |
| 100 | 0.0362 | 0.0348 | [0.0332, 0.0393] | 0 | 0.1881 | 0.0084 | 0.0425 | [0.0046, 0.0121] | -0.1277 | 0.1965 |
| 150 | 0.0321 | 0.0278 | [0.0297, 0.0345] | 0 | 0.1604 | 0.0156 | 0.0315 | [0.0128, 0.0183] | -0.0935 | 0.1513 |
| 200 | 0.0309 | 0.0239 | [0.0288, 0.033] | 0 | 0.1618 | 0.0191 | 0.026 | [0.0168, 0.0214] | -0.0365 | 0.1547 |
| 300 | 0.0288 | 0.0181 | [0.0272, 0.0304] | 0 | 0.091 | 0.0212 | 0.0191 | [0.0195, 0.0229] | -0.0264 | 0.0855 |
| 400 | 0.0277 | 0.0151 | [0.0264, 0.029] | 0.0001 | 0.0953 | 0.0215 | 0.0158 | [0.0201, 0.0229] | -0.0182 | 0.0917 |
| 600 | 0.0264 | 0.0121 | [0.0253, 0.0275] | 0.0016 | 0.0709 | 0.0221 | 0.0126 | [0.021, 0.0232] | -0.007 | 0.0681 |
| 800 | 0.0261 | 0.0104 | [0.0251, 0.027] | 0.0026 | 0.0643 | 0.0228 | 0.0107 | [0.0219, 0.0238] | -0.0006 | 0.0615 |
| 1000 | 0.0258 | 0.0094 | [0.0249, 0.0266] | 0.0033 | 0.0614 | 0.0231 | 0.0097 | [0.0222, 0.0239] | 0.0009 | 0.0612 |
| 1300 | 0.0252 | 0.0081 | [0.0245, 0.0259] | 0.0063 | 0.0514 | 0.0231 | 0.0083 | [0.0224, 0.0238] | 0.0035 | 0.0524 |
| 1600 | 0.0249 | 0.0072 | [0.0243, 0.0255] | 0.0081 | 0.0483 | 0.0231 | 0.0074 | [0.0224, 0.0237] | 0.0059 | 0.0503 |
| 2000 | 0.0248 | 0.0066 | [0.0242, 0.0254] | 0.0078 | 0.0451 | 0.0233 | 0.0067 | [0.0227, 0.0239] | 0.007 | 0.0431 |
| 2400 | 0.0249 | 0.0062 | [0.0244, 0.0255] | 0.0086 | 0.0425 | 0.0236 | 0.0063 | [0.023, 0.0242] | 0.0076 | 0.0431 |
| 2900 | 0.0246 | 0.0055 | [0.0241, 0.0251] | 0.0107 | 0.0392 | 0.0235 | 0.0057 | [0.023, 0.024] | 0.0089 | 0.0395 |
| 3401 | 0.0245 | 0.0052 | [0.024, 0.0249] | 0.0115 | 0.0418 | 0.0234 | 0.0053 | [0.023, 0.0239] | 0.0093 | 0.0399 |

*Note*. Effect size estimates are shown for non-cross-validated analysis (non-CV) and for the test sets of cross-validated analysis (model generalization to unknown data). Presented statistics are summarized across k=500 bootstrapped resamples of the respective sample size. mOFC, medial orbitofrontal cortex; SD, standard deviation; CI, confidence interval.

Table S4. Effect size estimates for **thalamus** voxel with reference effect size of $\boldsymbol{R^2_p}$**=.016** across sample size and cross-validation.

| Sample size | $R^2_p$ non-CV | | | | | CV-based $R^2_p$ test | | | | |
|---|---|---|---|---|---|---|---|---|---|---|
| | Mean | SD | 95% CI | Min | Max | Mean | SD | 95% CI | Min | Max |
| 25 | 0.0687 | 0.0966 | [0.0602, 0.0772] | 0 | 0.7335 | -0.1681 | 0.3309 | [-0.1972, -0.139] | -3.1349 | 0.7054 |
| 35 | 0.0488 | 0.0728 | [0.0424, 0.0552] | 0 | 0.6408 | -0.0744 | 0.1349 | [-0.0862, -0.0625] | -0.7445 | 0.6716 |
| 50 | 0.0373 | 0.0516 | [0.0328, 0.0419] | 0 | 0.3661 | -0.0335 | 0.0786 | [-0.0404, -0.0266] | -0.3015 | 0.3019 |
| 70 | 0.0305 | 0.0387 | [0.0271, 0.0338] | 0 | 0.2434 | -0.0169 | 0.0561 | [-0.0218, -0.012] | -0.1967 | 0.2227 |
| 100 | 0.0273 | 0.0305 | [0.0246, 0.03] | 0 | 0.1539 | -0.0027 | 0.0408 | [-0.0063, 0.0009] | -0.1737 | 0.1772 |
| 150 | 0.0229 | 0.0227 | [0.0209, 0.0249] | 0 | 0.1668 | 0.0043 | 0.0273 | [0.0019, 0.0067] | -0.078 | 0.1625 |
| 200 | 0.0205 | 0.0184 | [0.0189, 0.0221] | 0 | 0.1162 | 0.0073 | 0.0214 | [0.0055, 0.0092] | -0.0547 | 0.1127 |
| 300 | 0.0184 | 0.0144 | [0.0171, 0.0197] | 0 | 0.0917 | 0.0099 | 0.0158 | [0.0086, 0.0113] | -0.0369 | 0.0891 |
| 400 | 0.018 | 0.013 | [0.0168, 0.0191] | 0 | 0.0768 | 0.0112 | 0.0139 | [0.01, 0.0124] | -0.0258 | 0.0656 |
| 600 | 0.0171 | 0.0106 | [0.0162, 0.0181] | 0 | 0.0602 | 0.0126 | 0.0111 | [0.0116, 0.0136] | -0.0152 | 0.0548 |
| 800 | 0.0167 | 0.0089 | [0.0159, 0.0175] | 0.0004 | 0.0479 | 0.0132 | 0.0091 | [0.0124, 0.014] | -0.0081 | 0.0441 |
| 1000 | 0.0164 | 0.008 | [0.0157, 0.0171] | 0.0004 | 0.0489 | 0.0134 | 0.0081 | [0.0127, 0.0141] | -0.0049 | 0.046 |
| 1300 | 0.0162 | 0.0067 | [0.0156, 0.0168] | 0.0018 | 0.0422 | 0.0138 | 0.0068 | [0.0132, 0.0144] | -0.0043 | 0.0438 |
| 1600 | 0.0161 | 0.0062 | [0.0156, 0.0166] | 0.0021 | 0.0385 | 0.0139 | 0.0061 | [0.0134, 0.0144] | -0.0022 | 0.0371 |
| 2000 | 0.0159 | 0.0055 | [0.0154, 0.0164] | 0.0029 | 0.0393 | 0.014 | 0.0055 | [0.0135, 0.0145] | -0.0001 | 0.0341 |
| 2400 | 0.0159 | 0.005 | [0.0154, 0.0163] | 0.0051 | 0.038 | 0.0142 | 0.005 | [0.0137, 0.0146] | 0.0006 | 0.0351 |
| 2900 | 0.0159 | 0.0045 | [0.0155, 0.0163] | 0.0062 | 0.0336 | 0.0143 | 0.0045 | [0.0139, 0.0147] | 0.0007 | 0.0312 |
| 3401 | 0.0157 | 0.0042 | [0.0154, 0.0161] | 0.0061 | 0.0314 | 0.0144 | 0.0042 | [0.014, 0.0147] | 0.0037 | 0.0287 |

*Note*. Effect size estimates are shown for non-cross-validated analysis (non-CV) and for the test sets of cross-validated analysis (model generalization to unknown data). Presented statistics are summarized across k=500 bootstrapped resamples of the respective sample size. mOFC, medial orbitofrontal cortex; SD, standard deviation; CI, confidence interval.

Table S5. Effect size estimates for **anterior mOFC** voxel with reference effect size of $\boldsymbol{R^2_p}$**=.005** across sample size and cross-validation.

| Sample size | $R^2_p$ non-CV | | | | | CV-based $R^2_p$ test | | | | |
|---|---|---|---|---|---|---|---|---|---|---|
| | Mean | SD | 95% CI | Min | Max | Mean | SD | 95% CI | Min | Max |
| 25 | 0.0597 | 0.0735 | [0.0532, 0.0662] | 0 | 0.4255 | -0.1678 | 0.3714 | [-0.2005, -0.1352] | -4.2755 | 0.4091 |
| 35 | 0.0358 | 0.0463 | [0.0318, 0.0399] | 0 | 0.309 | -0.0686 | 0.1103 | [-0.0783, -0.0589] | -0.7895 | 0.3473 |
| 50 | 0.0229 | 0.0324 | [0.0201, 0.0258] | 0 | 0.2114 | -0.0422 | 0.0739 | [-0.0487, -0.0357] | -0.5657 | 0.1821 |
| 70 | 0.0176 | 0.0227 | [0.0156, 0.0196] | 0 | 0.1553 | -0.0238 | 0.046 | [-0.0278, -0.0197] | -0.4222 | 0.1278 |
| 100 | 0.0142 | 0.0181 | [0.0126, 0.0158] | 0 | 0.1287 | -0.0113 | 0.0299 | [-0.0139, -0.0087] | -0.1407 | 0.1324 |
| 150 | 0.0115 | 0.0142 | [0.0103, 0.0128] | 0 | 0.0852 | -0.0042 | 0.0192 | [-0.0059, -0.0025] | -0.0635 | 0.0851 |
| 200 | 0.0096 | 0.0111 | [0.0086, 0.0106] | 0 | 0.072 | -0.0025 | 0.0141 | [-0.0038, -0.0013] | -0.0564 | 0.0691 |
| 300 | 0.0079 | 0.0086 | [0.0071, 0.0087] | 0 | 0.0595 | -0.0006 | 0.0106 | [-0.0015, 0.0004] | -0.0263 | 0.0518 |
| 400 | 0.007 | 0.007 | [0.0064, 0.0076] | 0 | 0.0384 | 0.0008 | 0.0081 | [0.0001, 0.0015] | -0.0217 | 0.0346 |
| 600 | 0.0061 | 0.0057 | [0.0056, 0.0066] | 0 | 0.0295 | 0.0019 | 0.0062 | [0.0013, 0.0024] | -0.0126 | 0.0291 |
| 800 | 0.0059 | 0.0052 | [0.0055, 0.0064] | 0 | 0.0332 | 0.0025 | 0.0057 | [0.002, 0.003] | -0.011 | 0.0314 |
| 1000 | 0.0056 | 0.0042 | [0.0052, 0.006] | 0 | 0.0263 | 0.0027 | 0.0046 | [0.0023, 0.0031] | -0.0069 | 0.0238 |
| 1300 | 0.0054 | 0.0038 | [0.0051, 0.0057] | 0 | 0.0226 | 0.0031 | 0.0042 | [0.0027, 0.0034] | -0.0086 | 0.0203 |
| 1600 | 0.0053 | 0.0034 | [0.005, 0.0056] | 0 | 0.0227 | 0.0033 | 0.0037 | [0.003, 0.0036] | -0.0055 | 0.0201 |
| 2000 | 0.0051 | 0.0029 | [0.0048, 0.0053] | 0 | 0.019 | 0.0034 | 0.0031 | [0.0032, 0.0037] | -0.0038 | 0.0188 |
| 2400 | 0.0051 | 0.0027 | [0.0049, 0.0054] | 0.0001 | 0.017 | 0.0037 | 0.0028 | [0.0034, 0.0039] | -0.003 | 0.0155 |
| 2900 | 0.0051 | 0.0025 | [0.0049, 0.0053] | 0.0003 | 0.015 | 0.0038 | 0.0025 | [0.0036, 0.0041] | -0.0018 | 0.0135 |
| 3401 | 0.005 | 0.0023 | [0.0048, 0.0052] | 0.0005 | 0.0132 | 0.0039 | 0.0024 | [0.0037, 0.0041] | -0.0011 | 0.0125 |

*Note*. Effect size estimates are shown for non-cross-validated analysis (non-CV) and for the test sets of cross-validated analysis (model generalization to unknown data). Presented statistics are summarized across k=500 bootstrapped resamples of the respective sample size. mOFC, medial orbitofrontal cortex; SD, standard deviation; CI, confidence interval.

Table S6. Effect size estimates for **Calcarine** voxel with reference effect size of $\boldsymbol{R^2_p}$**<.0001** across sample size and cross-validation.

| Sample size | $R^2_p$ non-CV | | | | | CV-based $R^2_p$ test | | | | |
|---|---|---|---|---|---|---|---|---|---|---|
| | Mean | SD | 95% CI | Min | Max | Mean | SD | 95% CI | Min | Max |
| 25 | 0.0558 | 0.0729 | [0.0494, 0.0622] | 0 | 0.4287 | -0.149 | 0.2586 | [-0.1717, -0.1263] | -1.8284 | 0.3996 |
| 35 | 0.0359 | 0.048 | [0.0316, 0.0401] | 0 | 0.2756 | -0.0702 | 0.1292 | [-0.0815, -0.0588] | -0.963 | 0.274 |
| 50 | 0.0237 | 0.0322 | [0.0209, 0.0265] | 0 | 0.2259 | -0.0373 | 0.0637 | [-0.0429, -0.0317] | -0.4153 | 0.2069 |
| 70 | 0.0164 | 0.02 | [0.0147, 0.0182] | 0 | 0.1293 | -0.0243 | 0.0394 | [-0.0278, -0.0209] | -0.2489 | 0.1127 |
| 100 | 0.0108 | 0.0139 | [0.0096, 0.012] | 0 | 0.0863 | -0.0161 | 0.0273 | [-0.0185, -0.0137] | -0.1794 | 0.0944 |
| 150 | 0.0072 | 0.0096 | [0.0064, 0.008] | 0 | 0.0749 | -0.0099 | 0.0169 | [-0.0114, -0.0084] | -0.0749 | 0.0613 |
| 200 | 0.0052 | 0.0075 | [0.0045, 0.0058] | 0 | 0.0576 | -0.0071 | 0.0119 | [-0.0081, -0.006] | -0.0543 | 0.0561 |
| 300 | 0.0033 | 0.0051 | [0.0028, 0.0037] | 0 | 0.0391 | -0.0043 | 0.008 | [-0.005, -0.0036] | -0.0348 | 0.0456 |
| 400 | 0.0026 | 0.0037 | [0.0023, 0.003] | 0 | 0.025 | -0.0029 | 0.0056 | [-0.0034, -0.0024] | -0.0228 | 0.0296 |
| 600 | 0.0017 | 0.0024 | [0.0015, 0.002] | 0 | 0.0186 | -0.0021 | 0.0038 | [-0.0024, -0.0017] | -0.0155 | 0.0182 |
| 800 | 0.0013 | 0.0018 | [0.0012, 0.0015] | 0 | 0.0127 | -0.0015 | 0.0027 | [-0.0018, -0.0013] | -0.0111 | 0.0123 |
| 1000 | 0.001 | 0.0015 | [0.0009, 0.0012] | 0 | 0.0115 | -0.0013 | 0.0022 | [-0.0015, -0.0011] | -0.0084 | 0.0117 |
| 1300 | 0.0008 | 0.0013 | [0.0006, 0.0009] | 0 | 0.0111 | -0.001 | 0.0017 | [-0.0011, -0.0008] | -0.0079 | 0.0099 |
| 1600 | 0.0007 | 0.0012 | [0.0006, 0.0008] | 0 | 0.0113 | -0.0007 | 0.0016 | [-0.0009, -0.0006] | -0.0066 | 0.0115 |
| 2000 | 0.0005 | 0.0009 | [0.0005, 0.0006] | 0 | 0.0076 | -0.0006 | 0.0012 | [-0.0007, -0.0005] | -0.0048 | 0.0069 |
| 2400 | 0.0005 | 0.0008 | [0.0004, 0.0005] | 0 | 0.0058 | -0.0005 | 0.001 | [-0.0006, -0.0004] | -0.004 | 0.0056 |
| 2900 | 0.0004 | 0.0006 | [0.0003, 0.0004] | 0 | 0.0054 | -0.0004 | 0.0008 | [-0.0005, -0.0004] | -0.0029 | 0.0054 |
| 3401 | 0.0003 | 0.0005 | [0.0003, 0.0004] | 0 | 0.0041 | -0.0004 | 0.0007 | [-0.0004, -0.0003] | -0.0028 | 0.004 |

*Note*. Effect size estimates are shown for non-cross-validated analysis (non-CV) and for the test sets of cross-validated analysis (model generalization to unknown data). Presented statistics are summarized across k=500 bootstrapped resamples of the respective sample size. mOFC, medial orbitofrontal cortex; SD, standard deviation; CI, confidence interval.

Figure S1. Non-thresholded brain-wide partial correlation between Body Mass Index and gray matter density in the full sample (n=3401) without cross-validation.

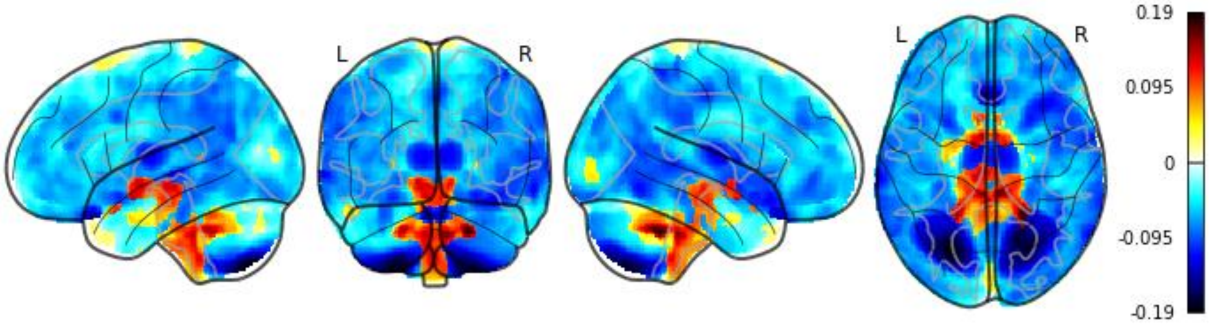


*Note. Glass brain projections are shown without applying a significance threshold. Color bar represents partial correlation coefficients calculated based on t-values from the two-sided contrast of the Body Mass Index predictor.*

Figure S2. Distribution of **partial *r*-**coefficients for **non-cross-validated** linear association between Body Mass Index and voxel-based morphometry across voxels and samples sizes.

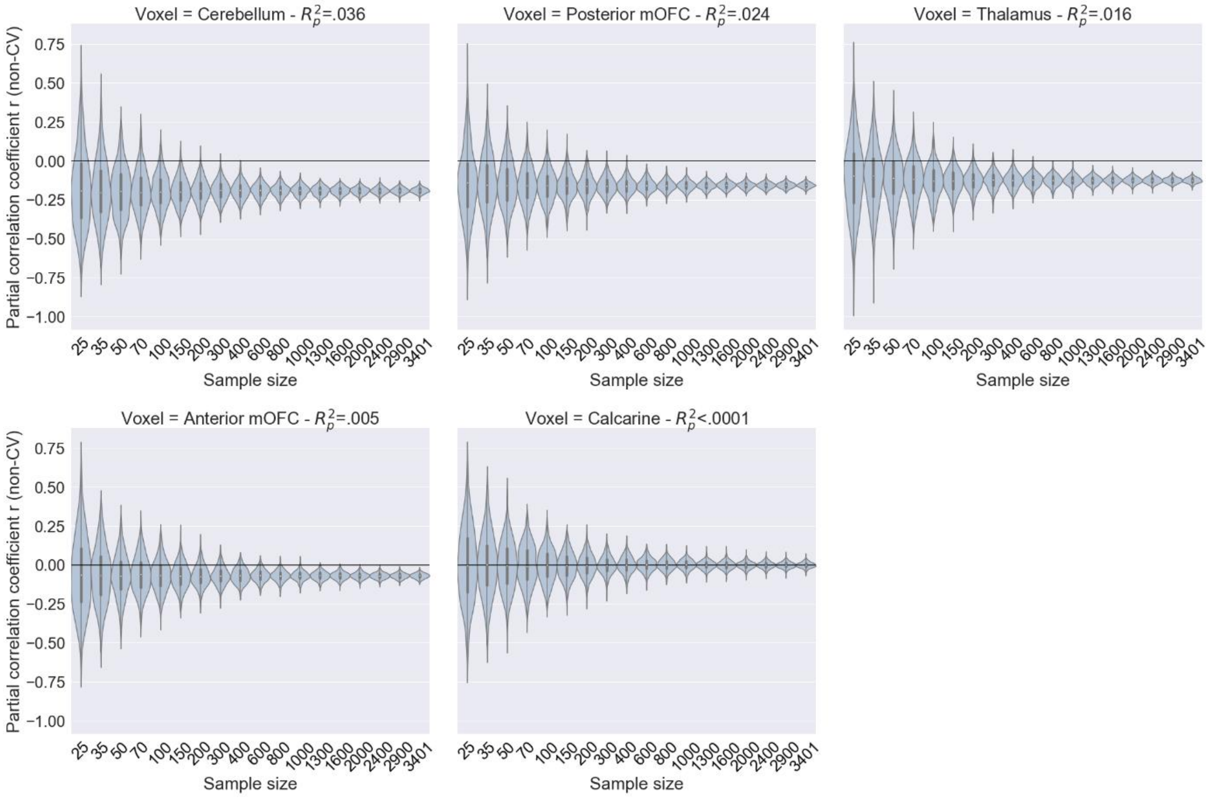


*Note. The distribution of effect size estimates is presented as a violin plot based on the resampling procedure.*

Figure S3. Distribution of **partial $R^2$** estimates for **non-cross-validated** linear association between Body Mass Index and voxel-based morphometry across voxels and samples sizes.

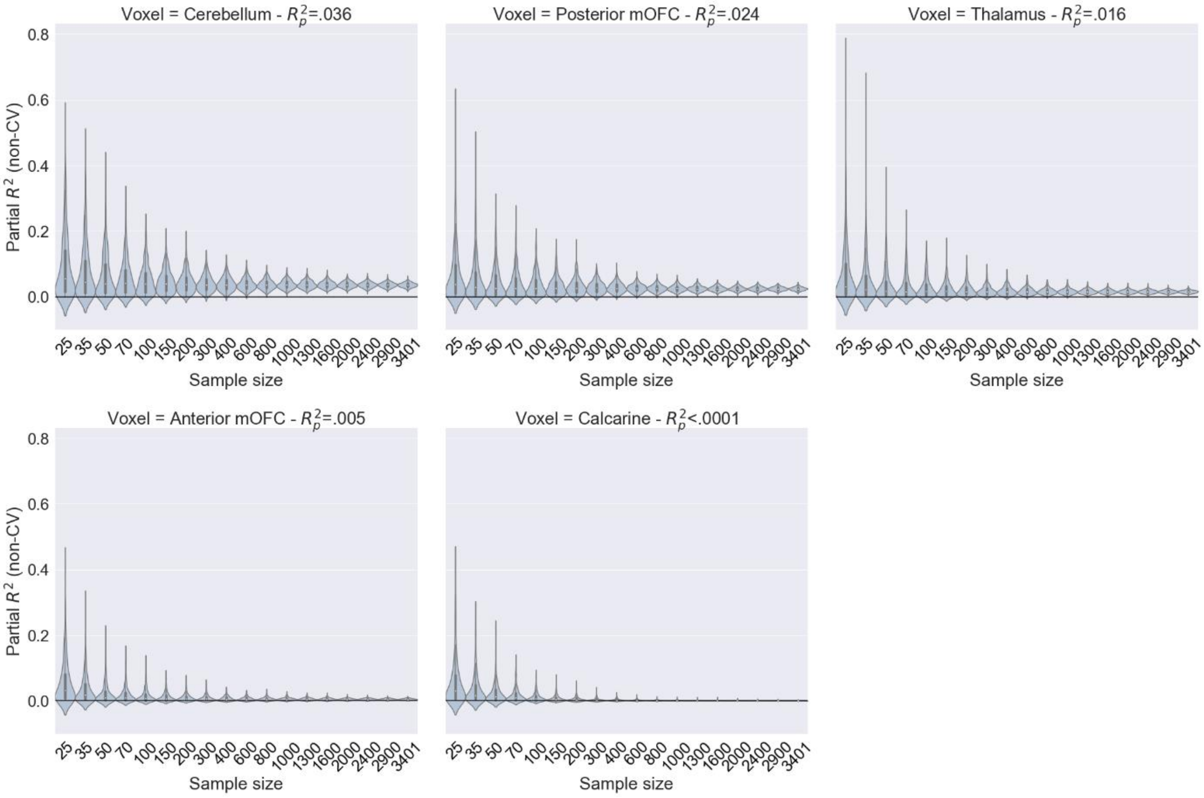


*Note. The distribution of effect size estimates is presented as a violin plot based on the resampling procedure.*

Figure S4. Distribution of **test-set partial *$R^2$*** estimates for **cross-validated** linear association between Body Mass Index and voxel-based morphometry across voxels and samples sizes.

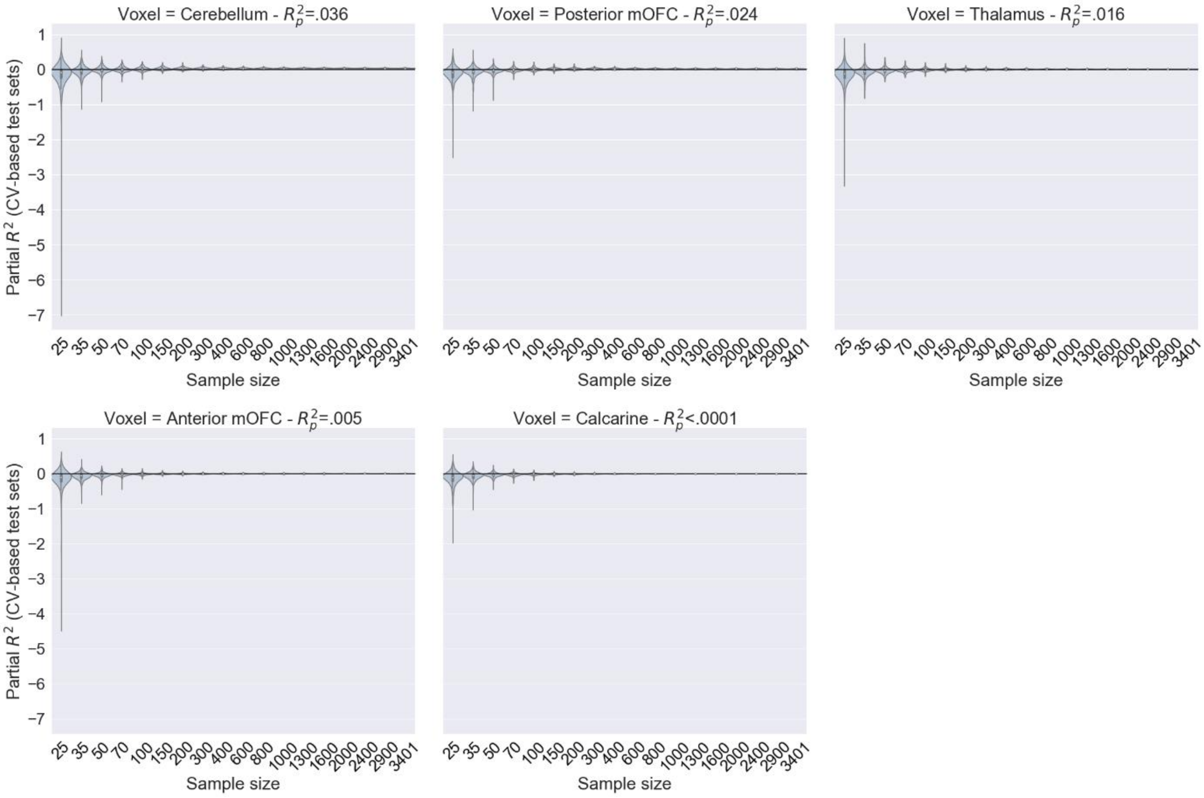


*Note. The distribution of effect size estimates is presented as a violin plot based on the resampling procedure.*

Figure S5. Association between non-cross-validated and cross-validation-based test set effect size estimates.

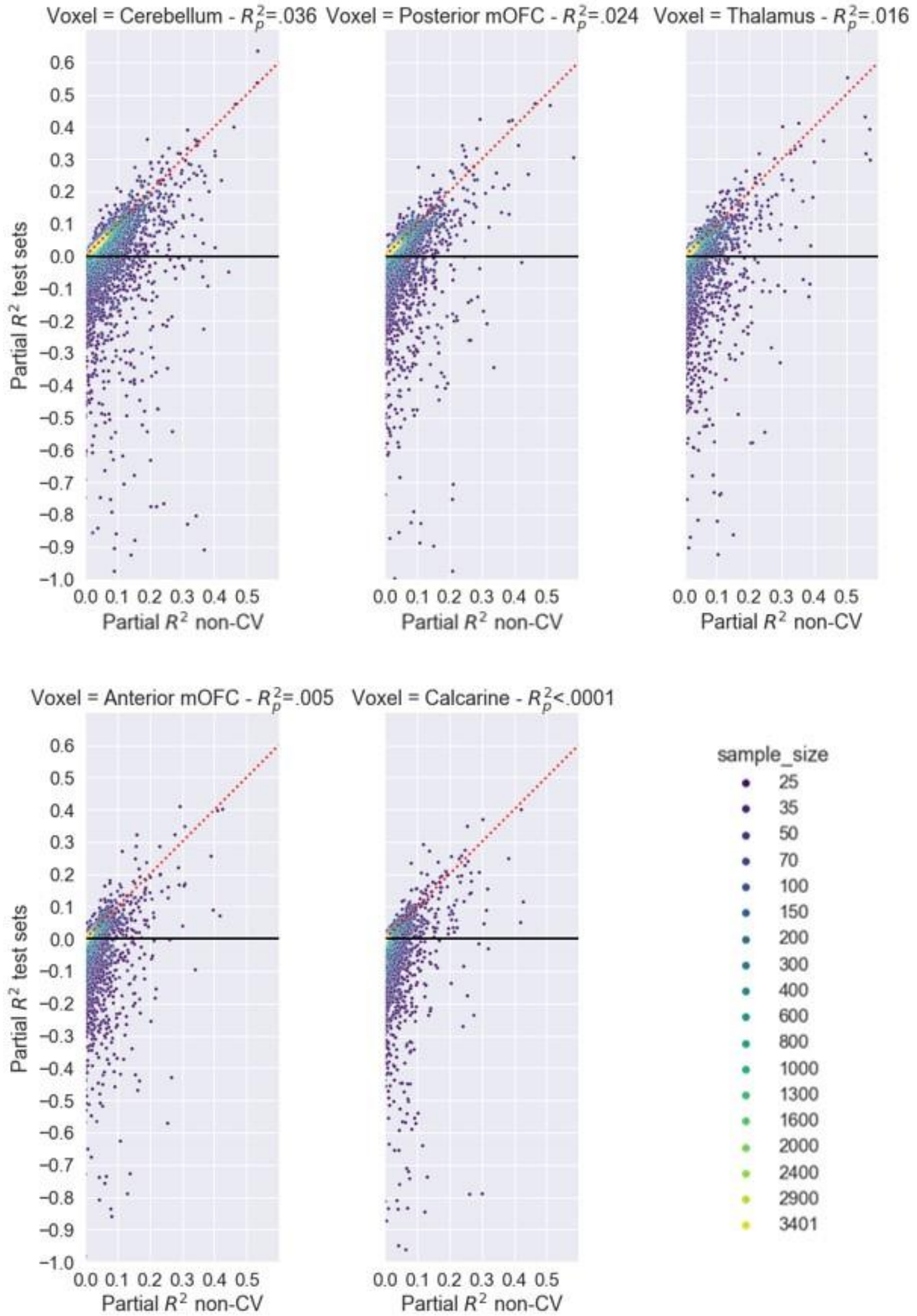


*Note. Effect size estimates are presented for each single run of the resampling procedure. The red dotted line represents an identity line: for data points below the line, the test set effect size is smaller than the non-cross-validated effect size for the specific sample and vice versa for data points above the line. Due to extreme outliers in the test set effect size estimates data are trimmed for better visualization – values below $R^2_p$=-1 are not shown. The color bar codes different samples sizes.*
